# The impact of regular school closure on seasonal influenza epidemics: a data-driven spatial transmission model for Belgium

**DOI:** 10.1101/230565

**Authors:** Giancarlo De Luca, Kim Van Kerckhove, Pietro Coletti, Chiara Poletto, Nathalie Bossuyt, Niel Hens, Vittoria Colizza

## Abstract

School closure is often considered as an option to mitigate influenza epidemics because of its potential to reduce transmission in children and then in the community. The policy is still however highly debated because of controversial evidence. Moreover, the specific mechanisms leading to mitigation are not clearly identified.

We introduced a stochastic spatial age-specific metapopulation model to assess the role of holiday-associated behavioral changes and how they affect seasonal influenza dynamics. The model is applied to Belgium, parameterized with country-specific data on social mixing and travel, and calibrated to the 2008/2009 influenza season. It includes behavioral changes occurring during weekend vs. weekday, and holiday vs. school-term. Several experimental scenarios are explored to identify the relevant social and behavioral mechanisms.

Stochastic numerical simulations show that holidays considerably delay the peak of the season and mitigate its impact. Changes in mixing patterns are responsible for the observed effects, whereas changes in travel behavior do not alter the epidemic. Weekends are important in slowing down the season by periodically dampening transmission. Christmas holidays have the largest impact on the epidemic, however later school breaks may help in reducing the epidemic size, stressing the importance of considering the full calendar. An extension of the Christmas holiday of 1 week may further mitigate the epidemic.

Changes in the way individuals establish contacts during holidays are the key ingredient explaining the mitigating effect of regular school closure. Our findings highlight the need to quantify these changes in different demographic and epidemic contexts in order to provide accurate and reliable evaluations of closure effectiveness. They also suggest strategic policies in the distribution of holiday periods to minimize the epidemic impact.

## 1 Background

Children represent an epidemiological group of central importance for the transmission of influenza [76, 95, 5]. They often have a larger vulnerability to infections because of limited prior immunity, and they mix at school with high contact rates [80] thus representing key drivers for influenza spread. The closure of school has been associated to the potential of reducing influenza propagation in the community by breaking important chains of transmission. It is expected to potentially delay the peak, and reduce the epidemic impact, at peak time and of the overall wave. Though not specifically recommended by the World Health Organization during the 2009 H1N1 pandemic, it is envisioned as a possible non-pharmaceutical intervention for pandemic mitigation left to the decision of national and local authorities [12, 98].

A large body of literature exists on the topic, however contrasting evidence lead to no definitive emerging consensus [22, 69]. Benefits and limitations appear to depend on the specific epidemic context. For example, influenza epidemics characterized by a larger attack rate in children compared to adults are expected to be more sensitive to the closure of schools [24]. This experience was reported in many countries during the 2009 H1N1 influenza pandemic, where closed schools coincided with a marked reduction of influenza activity [71, 82, 25, 64, 99, 28, 29, 79, 42, 43, 4, 33, 65]. School closure interventions are often considered along with other mitigation strategies, as it happened with social distancing in Mexico following the pandemic outbreak [35, 28], making their effect more difficult to isolate. Studies generally report a slowing down effect in the incidence during closure, however in some cases the effect may be mixed with the natural decline of the epidemic, because of late implementation [69]. In addition, no clear trend is observed in the impact of school closure on the epidemic burden depending on the time of closure – before, around, or after the peak [69].

Given its potentially important role in reducing the epidemic impact, school closure has more often been investigated in the realm of pandemics compared to seasonal influenza [69]. For the latter, regular school closure during holidays in temperate regions has been considered as a natural example to evaluate the impact of school closure [62, 23, 52]. Directly extending the application to pandemic situations has however important limitations. Closure associated to winter holidays is regularly scheduled in the school calendar, whereas school closure as an intervention corresponds to an unplanned interruption of school attendance that may take different forms (e.g. proactive vs. reactive, at the national or local level, with a gradual closure of classes or of the entire school) [22, 53, 49, 30]. In addition to the different nature of closure, also its duration changes from a fixed scheduled period of holidays to one of variable extension depending on the ongoing epidemic and resulting outcome. How individual behavior changes in all these conditions is the critical aspect to quantify in order to accurately assess the impact of school closure on transmission.

Transmission models fitting the 2009 H1N1 pandemic or parameterized to a similar pandemic scenario have been used to assess the value of school closure or summer holidays [79, 39, 4, 49, 30]. Few of them are based on estimates for social mixing changes [39, 4], as data collected during a pandemic are limited [68, 39], leaving other approaches to rely on assumptions about contacts that may critically affect the studies’ findings. Applications to seasonal influenza may on the other hand count on a more accurate description of population mixing. Surveys conducted over the calendar year to measure variations of mixing patterns [61, 41, 40, 13] offer indeed the opportunity to perform data-driven modeling studies that mechanistically assess the role of school holidays on seasonal influenza. Interestingly, they also highlighted considerably large differences across countries in the way contacts change from term-time to school holidays [61], suggesting the need for country-specific estimates to accurately and reliably parameterize models.

Changes in mobility is another important aspect that is rarely integrated in school closure studies. Travel is known to be responsible for the spatial dissemination of influenza [57, 94, 32, 46, 44, 47, 9, 10, 7, 4]. In addition to extraordinary travel drops in reaction to epidemics [2, 7, 84], mobility changes regularly occur during school holidays compared to term-time [75, 45]. Moreover, important differences were highlighted in the mobility of children vs. adults and their associated variations, so that their coupling with social mixing changes occurring during holidays may have a considerable impact on the epidemic outcome [4, 45].

Our aim is to explicitly integrate social mixing and travel from data into a modeling framework to assess how variations induced by regular school closure may impact seasonal influenza epidemics. Three modeling studies were developed so far with similar objectives. Towers and Chowell studied the impact of day-of-week variations in human social contact patterns on incidence data collected at a large hospital in Santiago, Chile, during 2009 H1N1pdm [90]. Their approach was not spatial, therefore did not consider mobility changes, and mainly focused on the sensitivity of influenza incidence variations to the latency period. Apolloni *et al*. used a stylized analytical approach to evaluate the role of age-dependent social mixing and travel behavior on the conditions for epidemic spatial invasion [4]. The model can compare different contexts, with or without schools in terms, and also account for associated changes. The contexts are however considered independently (no full school calendar can be considered) and the epidemic impact is evaluated only in terms of conditions for spatial dissemination. Going beyond these limitations, more recently Ewing *et al*. introduced an age-specific spatial metapopulation model to evaluate how behavioral changes associated to winter holiday impact the flu season [45]. The model is applied to the United States and it integrates data on travel behavior, whereas mixing is assumed from estimates available from Europe and adapted to summer holiday changes measured in the UK during the 2009 pandemic [39]. Their findings identify changes in mixing patterns as the key element responsible for the epidemic effects induced by holidays.

Given the central role of mixing patterns largely supported by evidence [22, 69, 4, 45], the heterogeneous country-specific contact variations measured in Europe [80, 61], and the marked difference expected in individual behavior during a seasonal flu epidemic vs. a pandemic, here we extend prior approaches to introduce a data-driven spatially explicit model fully parameterized on Belgium. The aim is to reduce assumptions in favor of input data, and to exclusively focus on seasonal influenza and associated parameterization. Contact data associated to four types of calendar days are considered, belonging to Regular Weekday, Regular Weekend, Holiday Weekday, Holiday Weekend (here ‘regular’ refers to non-holiday period), allowing us to assess the role of weekends in addition to holidays. A richer calendar with additional holidays beyond Christmas break is also considered. Confirming and extending prior results with a different modeling approach and input data would greatly support our understanding of the role of changes in mixing and travel on influenza epidemics.

## 2 Methods

In order to study the role of changes in contact patterns and in travel behavior along the calendar, we considered a mathematical approach for the spatial transmission of influenza in Belgium. We built a discrete stochastic age-specific spatial metapopulation model at the municipality level, based on demographic, mixing, and mobility data of Belgium. We parameterized it with influenza-like-illness (ILI) data reported by the Belgian Scientific Institute of Public Health at the district level for the 2008/2009 season. By using the Belgian school calendar for that season, we assessed the impact of the individual changes in mixing and travel behavior during regular school closure, given by available data. We then tested experimental scenarios to identify the mechanisms responsible for the observed epidemic outcomes. Here, we describe in detail the mathematical model, input data, calibration procedure and experimental scenarios.

### 2.1 Age-specific metapopulation model

Metapopulation epidemic models are used to describe the spatial spread of an infectious disease through a spatially structured host population [59, 78, 58, 73]. They are composed of patches or subpopulations of the system, connected through a coupling process generally describing hosts mobility. Here we consider the population to be divided into two age classes, children and adults, based on the modeling framework introduced by Apolloni et al. [4]. Infection dynamics occur inside each patch, driven by the contacts between and within these two classes, and spatial spread occurs via the mobility of individuals (Figure 1). Both processes are modeled explicitly with a discrete and stochastic approach. The model is based on Belgian data and follows the time evolution of the 2008/2009 school calendar. It includes 589 patches corresponding to the 589 municipalities (nl. *gemeenten*, fr. *communes*) of Belgium. Weekends and school holidays are explicitly considered, and variations in mixing and travel behavior are accounted for in the model and based on data. In the following, we describe in detail the various components of the model.

#### 2.1.1 Demography and social mixing

Individuals are divided into children (*c*, age less than 19y) and adults (*a*, otherwise). Population size and age structure per municipality as of January 1, 2008 are obtained from Belgian Statistics [1].

Social mixing between the two age groups is quantified by contact matrices extracted from the data obtained through a Belgian social contact survey [80, 61]:

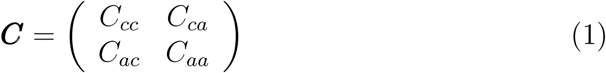

where the element *ij* (*i* = *a, c*, *j* = *a, c*) is given by:

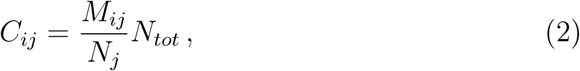

with *M_ij_* the average number of contacts made by survey participants in age class *i* with individuals in age class *j*, *N_j_* the population of age class *j*, *N_tot_* the total population of Belgium. ***C*** is defined at the national level, and here we assume that it is the same throughout the country, with the number of contacts being altered exclusively by the patch demography. From survey data, we have that the contact matrix for a regular weekday is:

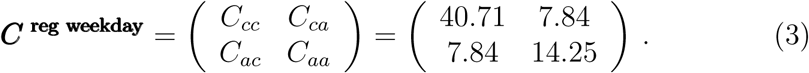

The variations for the other day types are discussed in paragraph 2.1.4.

**Figure 1:**
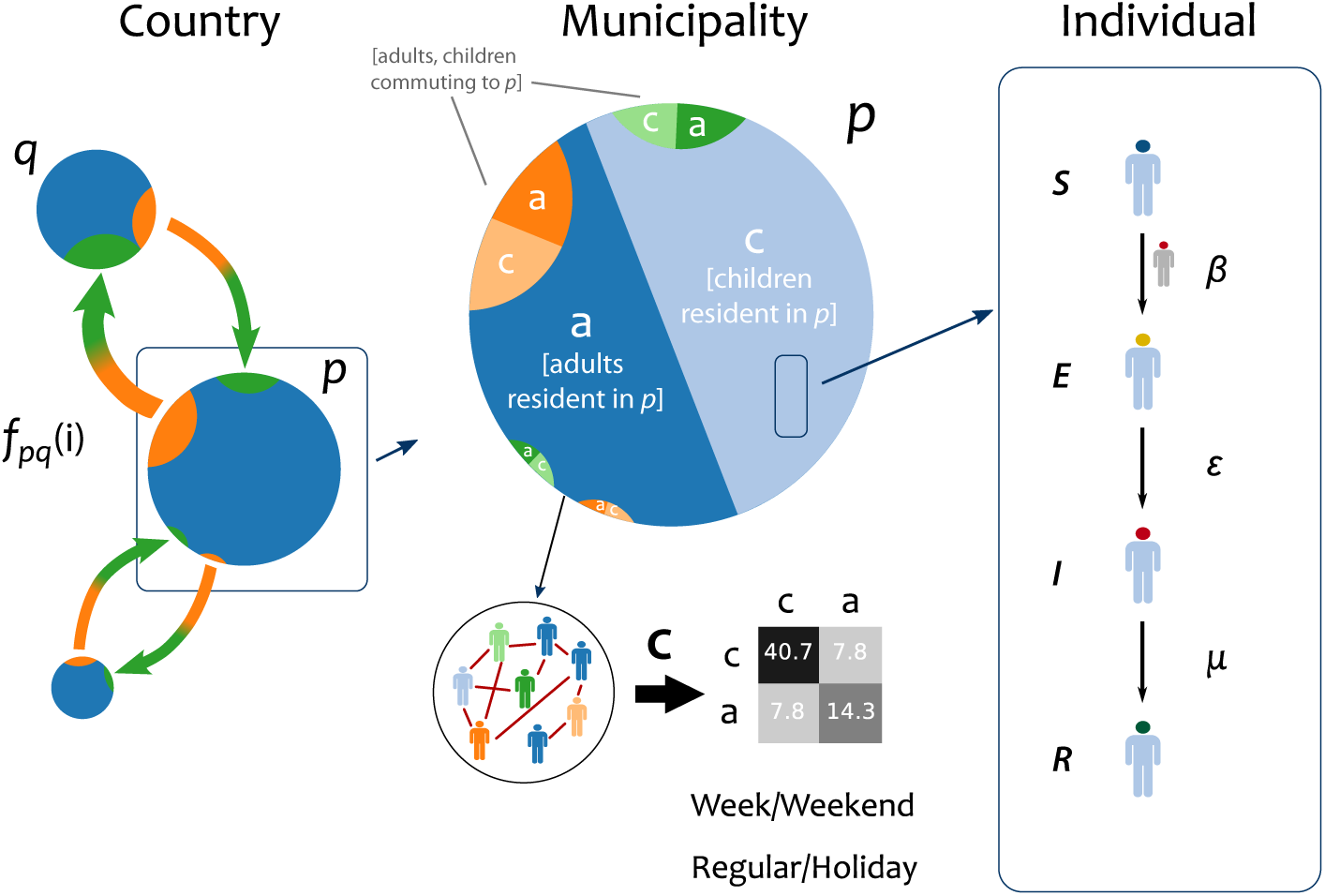
Schematic illustration of the spatial age-structured metapopulation model. The metapopulation modeling scheme is composed of three layers. At the country scale, Belgium is modeled as a set of patches (here indicated with *q* and *p*) corresponding to municipalities coupled through mobility of individuals *f_pq_*(*i*) of age class *i* at time *t*. Within each municipality, population is divided into two age classes, children (*c*) and adults (*a*), whose mixing pattern is defined by the contact matrix ***C***. Individuals resident of patch *p* and individuals commuting to that patch (e.g. resident of patch *q*) mix together following commuting. The figure reports as an example the contact matrix of a regular weekday (Eq. (3)). Mobility and mixing vary based on the calendar day (regular/holiday, weekday/weekend). Influenza disease progression at the individual level is modeled through a Susceptible-Exposed-Infectious-Recovered compartmental scheme, with *β* indicating the per-contact transmission rate, *∊* the rate from exposed to infectious state, *μ* the recovery rate.

#### 2.1.2 Infection dynamics

Influenza disease progression is described through a Susceptible-Exposed-Infectious-Recovered (SEIR) model (Figure 1) [3, 74]. A susceptible individual can contract the disease with a per-contact transmissibility rate *β* from infectious individuals, then entering the exposed or latent class. After an average latency period of *∈*^−1^ = 1.1 days [15, 20], the individual becomes infectious for an average duration *μ*^−1^ = 3 days [15, 20] and can transmit the infection, before recovering and becoming immune to the disease. A fraction of children (*g_c_*) and adults (*g_a_*) were considered immune to the disease at the beginning of the influenza epidemic, based on available knowledge on prior immunity and vaccination coverage in the country for the 2008/2009 influenza season and prior seasons (*g_c_* = 39.87%, *g_a_* = 53.19%) [100, 72, 89].

The force of infection for a susceptible individual of age class *i* (*i* = *a*, *c*) in a given patch *p* is given by:

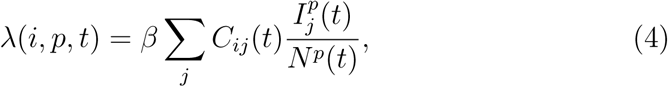

where *j* runs on age classes, 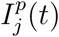 and *N^p^*(*t*) count the total number of infectious individuals of age class *j* and the total population size of patch *p* at time *t*, respectively. This quantity changes with time for two reasons. The first is the school calendar, distinguishing between regular weekdays, regular weekends, holiday weekdays, holidays weekends, and accounted for by *C_ij_*(*t*). The second is the mobility of individuals. At a given day of the simulation, each patch *p* may include: non-commuting residents of *p*, commuters from neighboring patches for school/work, commuting residents of *p* after school/work (see paragraph 2.1.3).

#### 2.1.3 Mobility

Coupling from patch *p* to patch *q* is given by the mobility of age class *i*, i.e. *f_pq_*(*i*) (Figure 1). We considered commuting data across patches from the *2001 Socio Economic Survey* of the Belgian Census [66] to describe the regular mobility of individuals for school/work during a regular weekday. Data are not age-specific, so we extracted the commuting fluxes per age class based on the probability of children (adults) of commuting on a given distance computed on the French commuting data [67]. Such inference was based on the assumption of a similar mobility behavior across the two neighboring countries.

Air travel was not considered due to negligible internal air traffic within the country.

#### 2.1.4 Changes in social mixing and travel behavior during school closure

Changes in social mixing are based on the data of the Belgian social contact survey [80, 61], where participants were asked to report their number of contacts during a regular weekday, a regular weekend, a holiday weekday or a holiday weekend. In addition to the contact matrix of Eq. (3), we have:

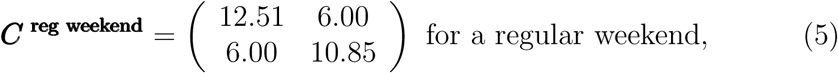

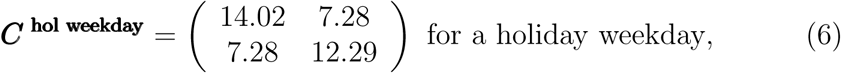

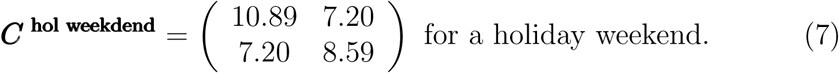

Concerning variations in mobility, schools are closed during weekends and holidays, so no commuting exists for children in those days. We considered adults to continue commuting during holiday weekdays, assuming that adults’ time off of work would be homogeneously distributed throughout the year. Concerning adult mobility during the weekends, we estimated the travel fluxes reductions based on statistics available for France [88], based on the same assumptions explained in paragraph 2.1.3. The resulting age-specific reductions for mobility are defined in Table 1.

**Table 1:** Age-specific mobility reductions

|  | children | adults |
| --- | --- | --- |
| Regular weekday | 1 | 1 |
| Holiday weekday | 0 | 1 |
| Saturday | 0 | 0.4053 |
| Sunday | 0 | 0.1473 |

### 2.2 Numerical simulations

Time is discretized considering a time step of *dt* = 0.5 days, with one timestep corresponding to the activities performed during a workday (i.e. commuting, social mixing), followed by a time step corresponding to the activities performed out of that timeframe (i.e. social mixing), as typically done in agent-based epidemic models [79]. Influenza transmission within each patch is modeled with binomial processes. Starting from the initial conditions set by influenza-like-illness surveillance data, we performed 2 · 10^3^ stochastic runs for each model under study.

### 2.3 Belgian school calendar for 2008/2009

Our model was based on the official Belgian school calendar for school year 2008/2009. Classes in Belgium are in session from Monday to Friday, and schools are closed during the weekends. The calendar included the following holidays for the 2008/2009 academic year during which schools were closed:

- Fall holiday: from October 25 to November 2, including the public holiday of the first and second of November;
- Public holiday of November 11;
- Christmas holiday: from December 20, 2008 to January 4, 2009;
- Winter holiday: from February 21 to March 1;
- Easter holiday: from April 4 to April 17;
- Long weekend: from May 1 to May 3, around the public holiday of May 1;
- Long weekend: from May 21 to May 24, around the public holiday of May 21;
- Long weekend: from May 30 to June 1, around the public holiday of June 1.

From July 1 to August 31 schools are closed for the summer holidays.

### 2.4 Influenza surveillance data for 2008/2009 season

We used influenza surveillance data collected by the Belgian Scientific Institute of Public Health [91]. Data report new ILI episodes registered each week by the network of sentinel general practitioners (GP). ILI is defined as sudden onset of symptoms, high fever, respiratory symptoms (cough, sore throat) and systemic symptoms (headache, muscular pain). For every episode, additional information is reported: age group (<5, 5-14, 15-64, 65-84, 85+), hospitalization, antiviral treatment, vaccination status, municipality of residence. The use of ILI surveillance data to approximate influenza incidence is a usual practice [81, 96, 56] and in the case of Belgium previous work showed good agreement of ILI data with virological data [16] and robustness across different surveillance systems [92].

Surveillance data on the new number of cases were aggregated at the district level to reduce signal noise.

### 2.5 Calibration

The metapopulation model was calibrated to the 2008/2009 influenza season. Though the simulated dynamics is spatially explicit, calibration was performed on Brussels district only, i.e. by comparing the simulated incidence profile of Brussels to the incidence ILI data for that district. We did not consider calibrating the model also in the remaining districts, as these were used for validation.

The model was seeded with the first non-zero incidence value provided by surveillance data per district and accounted for possible sampling biases. We used a bootstap/particle filter Weighted Least Square (*WLS*) with 20 particles to calibrate our model fixing the epidemiological parameters described in paragraph 2.1.2 and obtain the per-contact transmissibility *β*. Calibration was performed on normalized incidence curves to discount effects due to unknown GP consultation rates.

### 2.6 Experimental scenario design

To assess the impact of variations in contacts and mobility due to school closures, we compared the *realistic* model based on the Belgian school calendar and integrating the changes described in paragraph 2.1.4 with a set of experimental scenarios that we describe here.

To estimate the relative importance of variations in social mixing vs. variations in travel behavior, we considered:

- the *travel changes* model, where only variations in mobility occurring during weekends and holidays are considered, whereas social mixing is fixed as on a regular weekday;
- the *mixing changes* model, where only variations in social mixing occurring during weekends and holidays are considered, whereas travel is fixed as on a regular weekday;
- the *regular weekday* model, where no variations are considered, and social mixing and travel behavior are fixed as in a regular weekday.

To assess the role of each school holiday period, we considered scenarios where each period was removed, one at a time:

- *w/o Fall holiday* model, where the holiday period from October 25 to November 2 was removed;
- the *w/o Christmas holiday* model, where the holiday period from December 20, 2008 to January 4, 2009 was removed;
- the *w/o Winter holiday* model, where the holiday period from February 21 to March 1 was removed;
- the *w/o Easter holiday* model, where the holiday period from April 4 to April 17 was removed.

In all cases, the holiday period is substituted by the regular course of the week, with regular weekdays and regular weekends. In addition, we tested the *w/o holiday* model, where all holiday periods of the calendar are removed, and only the week structure is kept. We also considered a synthetic scenario where we extended the Christmas holiday of one week, before the start of the break, or after its end, referred to as the *Christmas holiday extension* models.

To assess the interplay between the timing of the epidemic and that of the holiday periods, we considered anticipation and delays of the start of the epidemic season, as follows:

- the 4 weeks anticipation model (–4*w*), where the start of the simulated influenza epidemic is anticipated 4 weeks prior to the start of the *realistic* model calibrated on the empirical data;
- the 2 weeks anticipation model (–2*w*), as above with an anticipation of 2 weeks;
- the 2 weeks delay model (+2*w*), as above with a delay of 2 weeks;
- the 4 weeks delay model (+4*w*), as above with a delay of 4 weeks.

In all these cases, the start of the epidemic is the only aspect that is being altered, whereas the school calendar (and associated variations in social mixing and travel behavior) remains fixed.

### 2.7 Analyses

We analyzed the spatial distribution of the force of infection determined by the demographic profile in space. To do so, we studied the distribution of the patch reproductive number *R^p^* that can be calculated as the largest eigenvalue of the next-generation matrix 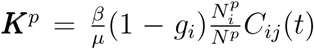 [38, 97]. This is done for the four types of days considered in terms of their variations of social mixing, namely regular weekday, regular weekend, holiday weekday, and holiday weekend.

Validation of the model is performed by comparing the simulated incidence profiles to the empirical surveillance data at the national and at the district level. In particular, we looked at the *peak difference* per district *d* per stochastic run *r*:

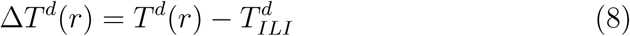

where *T^d^* (*r*) is the peak time of weekly incidence of run *r* in district *d* and 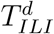 is the incidence peak reported from surveillance data in the same district. Medians per patch over 2 · 10^3^ stochastic runs are computed.

Scenario analyses are performed in order to assess the difference of an experimental scenario with the *realistic* model. We quantified the various comparisons in terms of:

- the *peak time difference* per patch 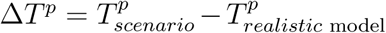, with *T^p^* the median peak time of the incidence curve in patch *p* computed on all stochastic runs (for both the scenario under study and the *realistic* model);
- the *peak incidence relative variation* per patch 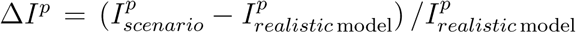, with *I^p^* the median incidence value at peak time in patch *p*;
- the *epidemic size relative variation* per patch 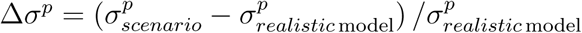, with *σ^p^* the median epidemic size in patch *p*.

Medians and 50%, 95% confidence intervals at the patch levels are computed for synthesis. In addition, medians, 50%, 95% confidence intervals of the simulated incidence ate also calculated at the national level across the tested scenarios.

## 3 Results

### 3.1 Reproducing the empirical influenza spreading pattern

Season 2008/2009 shows an ILI incidence that reaches its peak in week 5 of 2009, both in Brussels district and at the national level. The incidence is visibly slowed down during Christmas holiday (Figure 2), suggesting that holiday periods may have a measurable effect on transmission.

**Figure 2:**
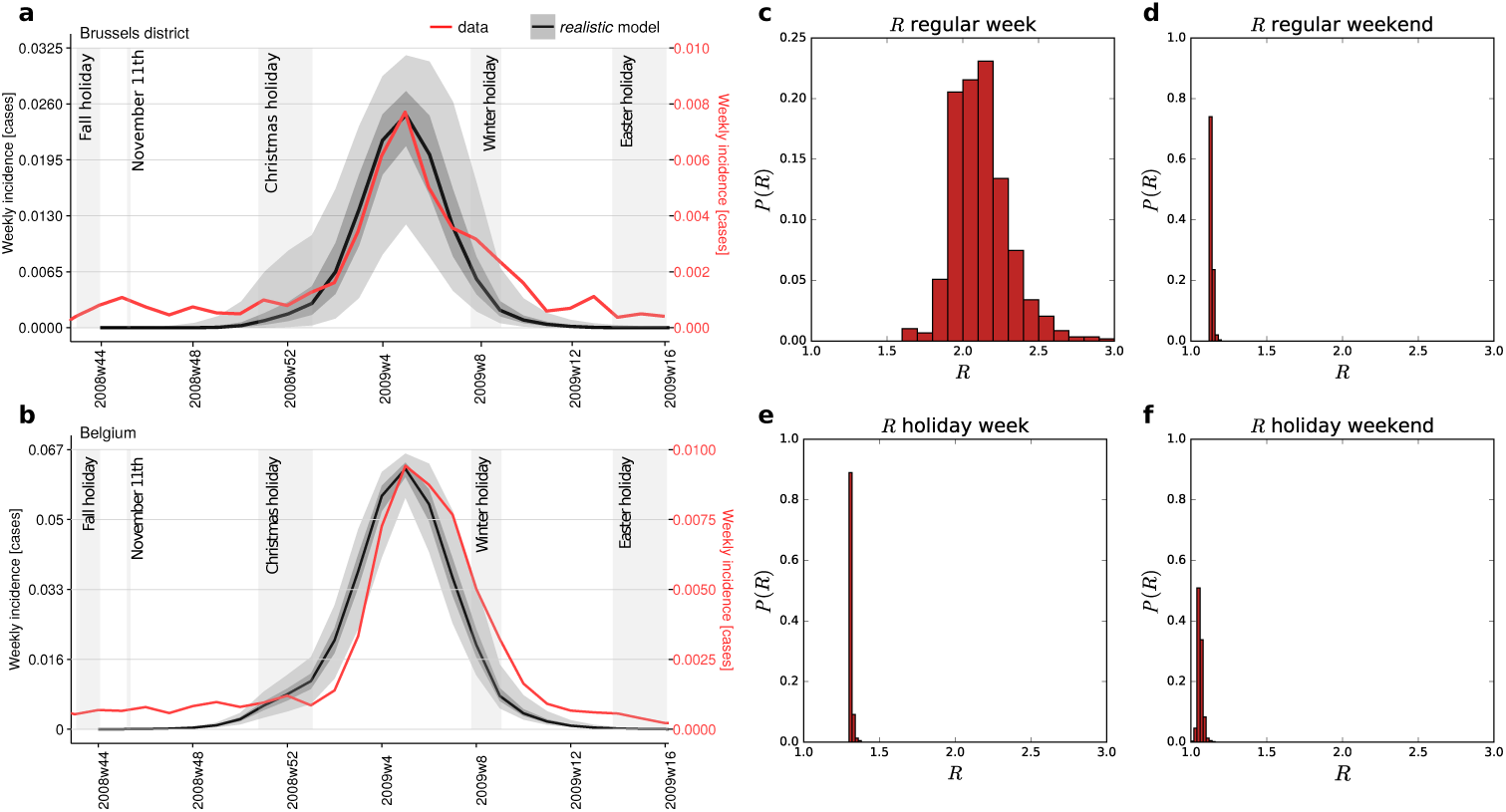
Calibration results. (a)-(b): Simulated and empirical incidence curves for the district of Brussels (panel a) and for the entire Belgium (panel b). The incidence curve of Brussels is the sole empirical data used for the calibration of the model. Different vertical axes referring to empirical (black curve, left axis) vs. simulated (red curve, right axis) incidences are used for the sake of comparison of the two curves. Different incidence values are due to unknown GP consultation rates characterising ILI surveillance data. (c)-(f): Probability distribution of the values of the reproductive number *R^p^* computed in each patch following the calibration. They refer to the different day types explored, i.e. belonging to a regular weekday (panel c), regular weekend (d), holiday weekday (e), holiday weekend (f).

The simulated peak time is found to be within one week of the empirically observed time for 76% of the districts, and within two weeks for 90% of them. Only two districts in the Province of Luxembourg showed greater discrepancies (four weeks). We observed a mild tendency towards a radial increase of the peak time difference from Brussels to the edge of the country, with 4 weeks difference obtained on the border between Belgium and Luxembourg.

The average patch reproductive number is estimated to be *R* = 2.12, corresponding to *β* = 0.0850 ([0.0674,0.0858] 95% confidence interval (CI)) of the per-contact transmissibility obtained from the calibration procedure (see Methods). The variation of *R^p^* at the patch level is given by the demographic profile of the population and its immunity profile. In addition, it also depends on the day type considered, whether regular or during a holiday, whether during the week or the weekend (Figure 2). Larger variations and higher values are obtained for a regular weekday, having the largest number of contacts, compared to less heterogeneous distributions and smaller *R^p^* values in the other cases. The patch reproductive number is lowest for the holiday weekend, corresponding to the lowest mixing.

### 3.2 Role of changes in individual behaviors during holidays and weekends: social mixing vs. travel

To assess the impact that changes in the social mixing or travel behaviors of individuals have on the epidemic outcome, we tested different experimental scenarios where we independently singled out these aspects. These scenarios are compared to the *realistic* model calibrated to the 2008/2009 influenza season, defined before, where all behavioral changes associated to the school calendar are considered.

Changes in individual behavior induced by weekends and holidays are found to strongly alter the epidemic dynamics leading to a considerable delay of the peak time (median of 3.7 weeks across patches, *regular weekday* model compared to the realistic model, Figure 3b) and smaller peak time incidence (33% median relative change, Figure 3d) and total epidemic size (11% median relative change, Figure 3c).

**Figure 3:**
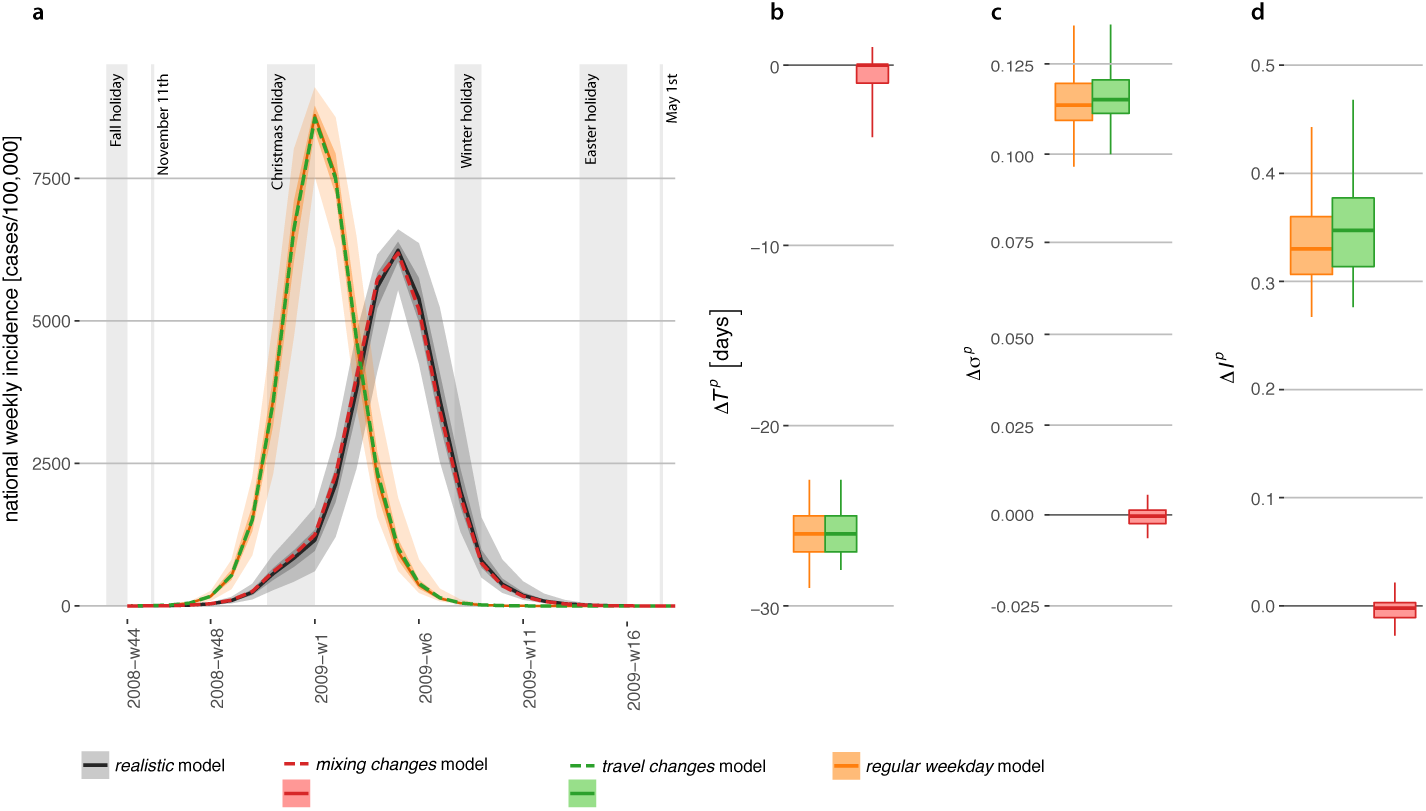
Role of social mixing vs. travel behavior. (a): Simulated weekly incidence profiles for influenza in Belgium. The *realistic* model is compared to the *travel changes* model, the *mixing changes* model, the *regular weekday* model. Median curves are shown for all cases, along with 50% confidence intervals (dark shade) and 95% CI (light shade), for the *realistic* and *regular weekday* model (they are not shown for the other models for the sake of visualization). (b)-(c)-(d): Peak time difference 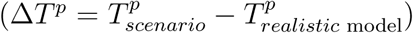, relative variation of epidemic size 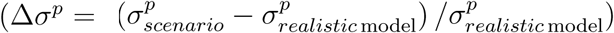 and relative variation of peak incidence 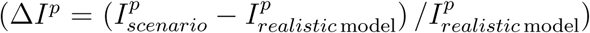, respectively, across the three experimental scenarios (see Methods for more details). Boxplots refer to the distributions across patches.

Once the variations in individual behavior affecting social mixing or mobility are considered in isolation, social mixing variation during weekends and holidays is found to be mainly responsible for the effects just described. The *travel changes* model is indeed comparable to the *regular weekday* model, whereas neglecting changes in mobility (*mixing changes* model) produces epidemic patterns very similar to the *realistic* model (zero median variations).

### 3.3 Role of distinct school holiday periods and possible holiday extensions

The school calendar in Belgium during the influenza season counts four long holiday periods: Fall holiday, Christmas holiday, Winter holiday, Easter holiday (see Methods).

Cumulatively, all holidays concur to delay the peak time of 1.7 weeks and to reduce the epidemic size of approximately 2%, with a reduction of the peak incidence (4%, all median values across patches, Figure 4). Among all holiday periods, the largest effect is produced by the Christmas holiday, responsible for the overall reduction of the epidemic size and a peak delay of about 1 week. The early break of Fall holiday has negligible impact instead. Winter holiday leads to a very small reduction of the epidemic size (median of 1%), but no effect on the peak timing or peak incidence. The impact of Easter holiday is negligible on all indicators. By comparing the effect of the *regular weekday* model (Figure 3) with the one of the *w/o holiday* model (Figure 4), both on the *realistic* model, we find that weekends have a major effect in slowing down the epidemic curve: a difference of Δ*T^p^* = −3.7, [−3.9, −3.6] weeks when no weekends are considered compared to Δ*T^p^* = −1.7, [−1.9, −1.2] weeks when they are included.

**Figure 4:**
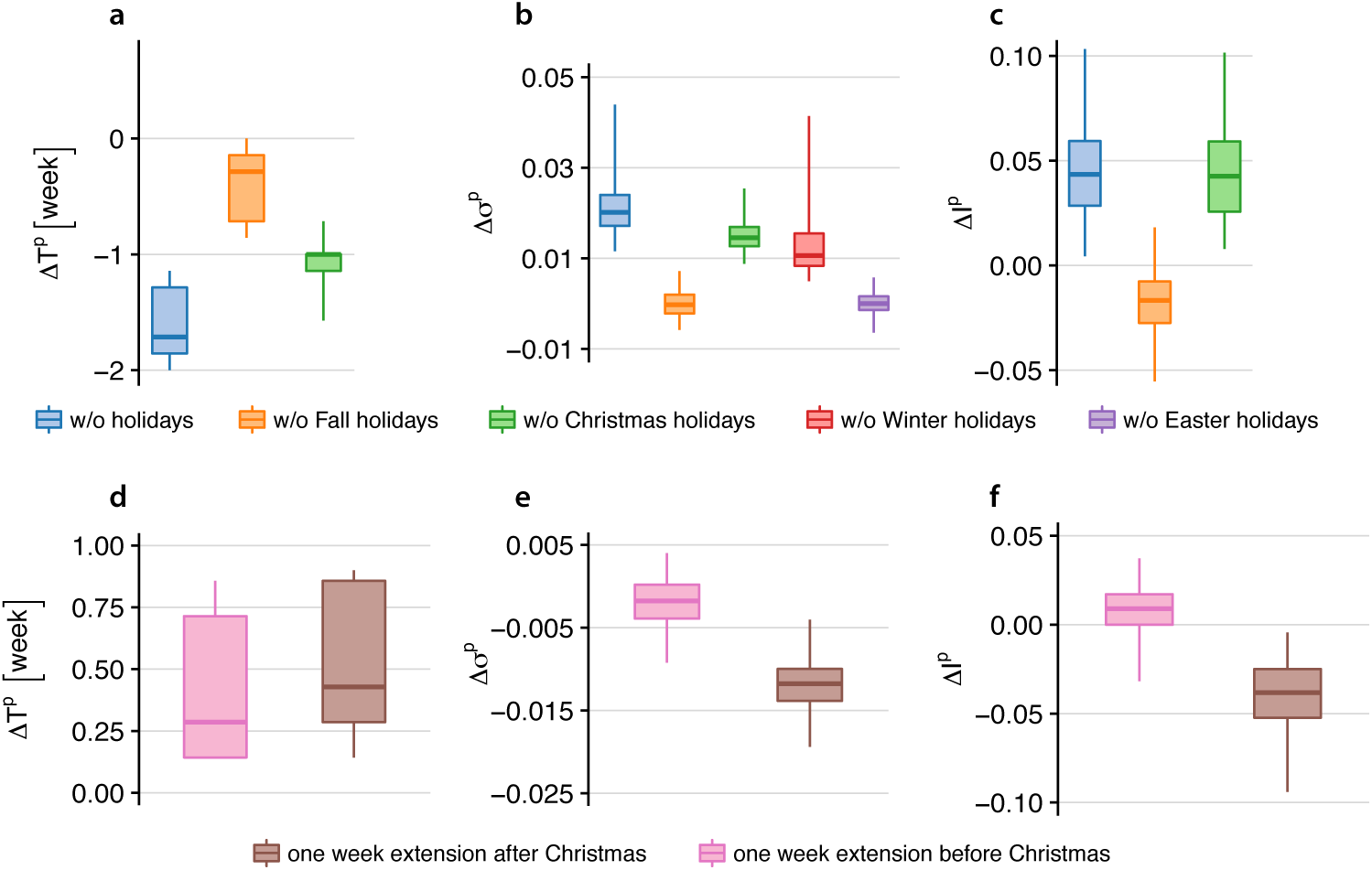
Impact of school holiday periods and holiday extensions. (a)-(b)-(c): Peak time difference, relative variation of epidemic size, and relative variation of peak incidence, respectively, across the following experimental scenarios: *w/o Fall holiday* model, *w/o Christmas holiday* model, *w/o Winter holiday* model, *w/o Easter holiday* model, *w/o holiday* model. Boxplots refer to the distributions across patches. (d)-(e)-(f): Peak time difference, relative variation of epidemic size, and relative variation of peak incidence, respectively, for the *Christmas holiday extension* models, before or after the break. Boxplots refer to the distributions across patches.

Given the major role of Christmas holiday, we also tested the effect of 1-week extension, before or after the break. The extension before Christmas holidays does not impact the resulting epidemic (Figure 4, panels d-e-f). If the additional week of holiday is considered after the break, no changes to the epidemic timing are observed, however the incidence at the peak decreases of 4% (median values), respectively.

### 3.4 Interplay of epidemic timing and school calendar: early vs. late influenza seasons

Christmas holidays are found to be the school closure period with the highest impact on the epidemic outcome, on both its timing and burden, for the 2008/2009 influenza season. Here we assess how this result may vary depending on the timing of the season, by investigating its interplay with the school closure calendar.

In order to distinguish between effects induced by the timing of the influenza season only and those related to other season-specific features (e.g. severity of the epidemic, strain circulation, weather, and others), we considered the same epidemic simulated with the *realistic* model. We explored anticipations and delays of this epidemic of two or four weeks and compared the results with the *realistic* model.

The strongest impact is observed for the earliest epidemic (–4*w* model) reporting a median anticipation of more than one week with respect to the realistic model (once discounted for the earlier start) and a median reduction of the peak incidence of about 10% (Figure 5). All other epidemics are rather similar to the realistic one, except for the – 2*w* model reporting a considerable reduction of the peak incidence (median of approximately 13% across patches). In addition, it is important to note that, differently from previous effects, the anticipation or delay of the season leads to a considerably larger variation of the simulated epidemic indicators across patches, signaled by the larger confidence intervals reported in Fig. 5.

**Figure 5:**
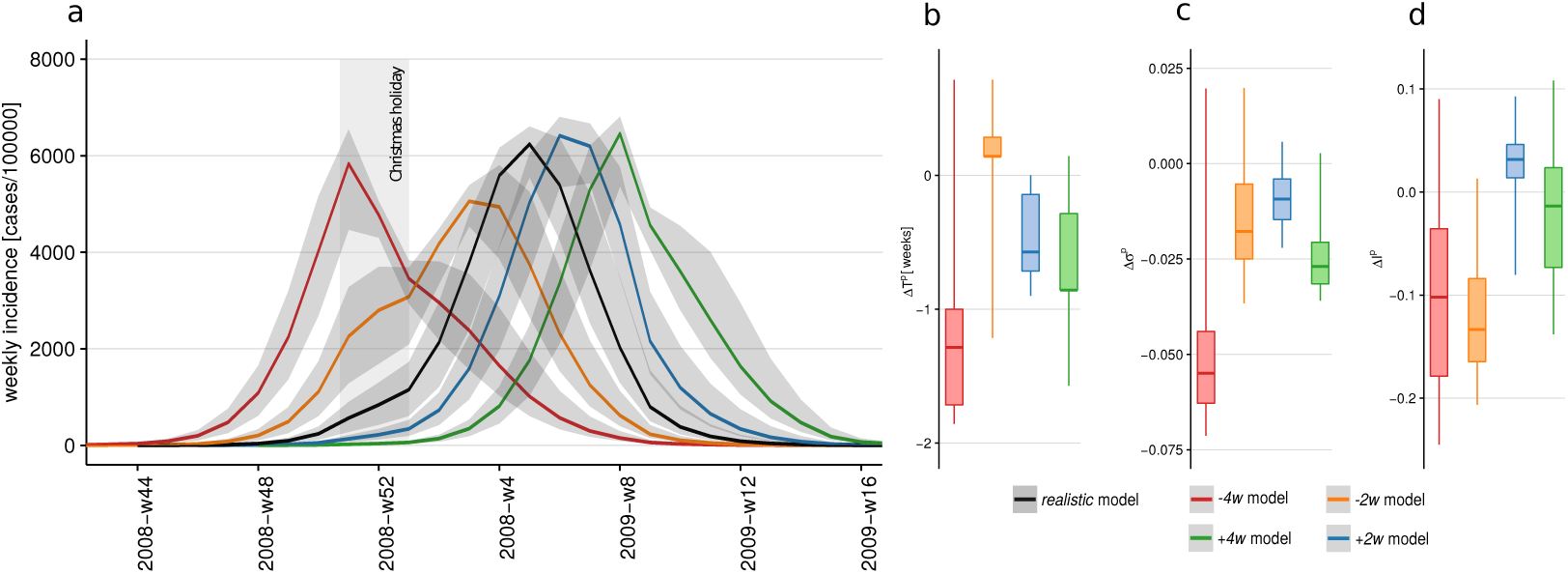
Effect of epidemic timing. (a): Simulated weekly incidence profiles for influenza in Belgium. The *realistic* model is compared to the scenarios considering the anticipation or delay of the epidemic (–4*w* model, –2*w* model, +2*w* model, +4*w* model). Median curves are shown along with 95% CI (light shade). (b)-(c)-(d): Peak time difference, relative variation of epidemic size and relative variation of peak incidence, respectively, across the considered experimental scenarios. Boxplots refer to the distributions across patches. The peak time difference Δ*T^p^* discounts the time shift of the initial conditions of the considered model.

## 4 Discussion

In this study, we considered the impact of regular school closure on the spatio-temporal spreading pattern of seasonal influenza. We focused on the case study of the 2008/2009 influenza season in Belgium. We used a spatial metapopulation model for the transmission of influenza in the country, based on data on contacts and mobility of individuals, and integrating data-driven changes in mixing and travel behavior during weekends and holiday periods.

The model calibrated on a single district (i.e. a subset of patches, ~ 3% of the country total) is able to reproduce with fairly good agreement the empirical pattern observed in the country for that season, suggesting that data-driven mixing and mobility are crucial ingredients to capture influenza spatial dynamics [94, 18, 36, 10, 79, 39, 4, 26, 45]. The result is a spatially heterogeneous propagation where the two ingredients act at different levels. Mixing is patch-dependent and determined by the local demography. The large variations observed in the distribution of children vs. adults lead to heterogeneous distributions of the values of the reproductive numbers per patch. In specific mixing conditions – e.g. those of a holiday weekend – a large fraction of patches has *R^p^* ≃ 1, indicating that those locations are found to be close to the critical conditions for epidemic extinction. Influenza is mainly sustained in patches having larger *R^p^* during those periods, and epidemic activity is then transferred to other patches through the mobility of infected individuals. Three-fourth of Belgian districts reach their epidemic peak in the simulations within one week of the empirical peak time. Districts exhibiting greater delays lie on the border of the country. This may be due to the model neglecting the mobility coupling between these regions and the neighboring countries, which is considerably large in some districts (e.g. the flux of individuals of the Luxembourg province commuting abroad represents almost 40% of the total flux of commuters of the district). Our study considered the country to be isolated for the sake of simplicity. We expect this border effect to be increasingly negligible for larger countries.

The simulated incidence profile clearly shows a slowing down in the growth of the number of new infections during the Christmas break, as reported by sentinel surveillance in the country, suggesting that holiday is associated to temporary reductions in influenza transmission. This was also found in previous empirical studies [23, 45]. To identify the mechanisms behind this effect, we isolated the changes in mixing and those in travel behavior during school closure, comparing different experimental scenarios, similar to Ewing et al. [45]. We found that mixing changes during weekends and holidays lead to a considerable delay of the epidemic, whereas travel changes would produce no noticeable effect. Moreover, changes in the social contacts would explain the entire difference observed between the *realistic* model based on the calendar and a model that does not include school closure. This confirms prior modeling findings on winter holidays for several influenza seasons in the United States [45]. In contrast to that work, we found however that an important mitigation of the epidemic impact at peak time also occurs, besides the peak delay.

The strong impact of the variation of mixing behavior is easily interpretable in terms of the reduction of the transmission potential expressed by the reproductive numbers per patch. This mainly results from the reduction of the number of contacts between children, as measured by the social contact survey conducted once schools are closed [61]. Travel changes, on the other hand, do not act directly on the transmission potential but affect the coupling force between epidemics in different patches and the opportunity for individuals to be exposed to the disease. For this reason, changes in travel behavior have a smaller effect that is found to be negligible for influenza epidemic spread, as also observed in the work of [45]. This is also consistent with the large literature on travel restrictions showing the little or no effect that these interventions have on pandemic spatial spread [32, 63, 46, 31, 44, 11, 18, 7, 84]. In addition to holidays, we also found that the closure of school during weekends has a visible effect on the epidemic, periodically dampening transmission, similarly to what observed in [90]. This is generally not reported by influenza surveillance systems, as data are collected on a weekly basis.

For the 2008/2009 influenza season we found that Christmas holiday, occurring during a growing phase of influenza activity, is the school break responsible for the largest impact in terms of timing (about 1 week anticipation if holiday is not observed), along with a 5% reduction of the epidemic size. If the school break occurs earlier (as for the Fall holiday) or much later in the influenza season (e.g. Easter holiday), no effect is produced on the resulting epidemic. The case of Winter holidays occurring during the fadeout of the epidemic shows that a small reduction of the total number of cases can still be achieved with school closure after the epidemic peak, whereas other studies showed minimal impact [45].

The analysis on a single season illustrates well how the epidemic impact of school closure depends on the interplay between closure timing and influenza season. By systematically exploring this interplay through synthetic scenarios, we confirm the importance of Christmas holiday in mitigating the influenza epidemic. Most importantly, we found that the break would have the largest impact for a very early season when school closing would occur at or around the epidemic peak. Reduction in transmission due to fewer contacts leads to a strong reduction of the incidence and ultimately of the total epidemic size, as also observed in pandemic settings [69]. In the other synthetic influenza seasons explored, a rebound effect was obtained when schools reopened after the break, most notably for the early season anticipating of two weeks the 2008/2009 influenza epidemic (-*2w* model). This was previously observed in other contexts [60, 48, 17, 8, 14, 62, 6, 35], also when no additional interventions beyond school closure were considered [48, 62]. The various tested scenarios show that Christmas break would have a larger mitigating impact if it occurs before (or around) the peak and when the incidence is about half the peak value or larger.

Our investigation shows that the role of holiday timing can be hardly inferred from few examples, and that other breaks beyond Christmas [45] may have an important mitigating impact. Also, the effect of a sequence of holidays occurring in an influenza season cannot be simply derived as a sum of the effects of each holiday period considered separately. Each break indeed affects the epidemic in a different way, altering its subsequent evolution in a non-linear way, so that the full calendar needs to be considered. Our findings help shedding light on previous empirical findings showing no clear pattern for the effects of school closure on peak incidence or total epidemic size, comparing closures before and after the peak [69].

In addition to school breaks already occurring in the calendar, we also explored a possible extension of Christmas holiday of one week. We chose this break as it led to the largest epidemic impact in our case study, and also because it generally occurs before the influenza epidemic peak (this is the case for all influenza seasons from the studied season to the current one, pandemic season excluded). As such, we expect it could have a favorable impact on the epidemic outcome in the majority of influenza seasons. Previous work analyzing the length of school closure found that two weeks or more appear to be enough to result in a recognizable effect [48, 62, 23, 99, 6], whereas shorter closures may not be beneficial or may not have an obvious impact [37, 34, 70, 86, 19]. Our synthetic results show that the extension would be advantageous only if implemented after the Christmas break, with a mitigation of the peak incidence and a minimal peak delay of a few days. Extensions are generally considered in the realm of reactive closures against a pandemic influenza. Here we decided to test this scenario as a regular closure given that – in a broader context – authorities in Belgium are currently discussing whether to modify the school calendar for pedagogical reasons: the aim would be to reduce summer holidays and redistribute holiday periods throughout the year [54]. We found that an extension of the Christmas holiday would be beneficial in the management of the influenza season potentially mitigating its epidemic impact.

Our findings are obtained on seasonal influenza, and results on peak delay were also recovered by modeling works on synthetic influenza pandemics considering reactive school closure [79]. The straightforward extension of our conclusions to the pandemic case faces however several challenges. First, effects induced by school closure may be specific to the specific epidemic profile, and therefore they may lead to different results depending on the pandemic under consideration [22]. For example, beneficial effects of school closure during 2009 H1N1 pandemic may have resulted from the larger magnitude of children attack rates vs. adults. Epidemic contexts more homogeneously impacting age classes may be less affected by school closure. Second, the nature of the school closure may alter the behavior of individuals during that period. In our study we considered holidays that are regularly planned in the school calendar and associated to specific social activities (e.g. vacation trips, family visits and others), for which contact data are available [61]. School closure during an influenza pandemic may be envisioned as a proactive or reactive measure to the ongoing outbreak. Not being planned, it is expected to have a stronger disruptive impact on social mixing of individuals on the short term compared to regular closure. On the other hand, it is argued that prolonged closure may limit the reduction of contacts on the long term, because of costs and logistics, and reduction in compliance rate [23, 22]. Having shown here that changes in social mixing represent the single element critically responsible for the impact of school closure on the epidemic outcome, we note that modeling results on school closure in the case of a pandemic would strongly be affected by assumptions considered for mixing changes, in absence of data.

While a large body of literature has recently focused on behavioral changes during an epidemic [51, 83, 85, 50, 93], still little is known to quantify them [68, 39, 61, 41, 40, 101, 102, 27, 77]. Our work focused on Belgium, as a rather detailed survey was conducted in the country to estimate contact rates in the population of different age classes at different periods of the calendar year [61]. These estimates constituted the input data to parameterize our spatial modeling framework. Modeling approaches to study epidemics in settings where no data exist are often based on the assumption that mixing would reduce following school closure and import estimates available from other settings or epidemiological contexts [79, 49, 45]. This may lead to several issues. Contacts and their changes along the calendar may be country-specific [80], thus affecting epidemic results when applied to a different context. Estimates of the overall reduction of the number of contacts during school closure vary widely. Transmission models fitted to epidemic data estimated reductions ranging from 16-18% for holidays during seasonal influenza in France [23], to 25% in Hong Kong for proactive school closure during the 2009 H1N1 pandemic [99], to 30% for the social distancing interventions (including school closure) implemented in Mexico following the start of 2009 pandemic [28]. A large-scale population-based prospective survey in Europe estimated the changes in contact patterns for holiday versus regular period to correspond to a reduction in the reproductive number as high as 33% for some countries, whereas for others no significant decrease was observed [61]. Finally, such overall reduction does not allow to fully parametrize a contact matrix. Such evidence does not support the parameterization of mixing changes from different countries and/or epidemic situations (e.g. seasonal vs. pandemic) [45]. The reduction is expected to be heterogeneous across mixing groups, because of compensatory behaviors (e.g. children drastically reduce children-children contacts but increase children-adults contacts during holidays) [61, 40]. Assumptions on the relative role of specific age classes in absence of data may lead to biases in the modeled epidemic outcome, especially for epidemics reporting large differences in attack rates in children vs. adults. Our work highlights the need to expand our knowledge on contacts and associated changes induced by social activity or by the epidemic itself, in order to better parameterize models and provide reliable and accurate results for epidemic management.

Our study has a set of limitations that we discuss in the following. The host population is divided into two classes only. While a larger heterogeneity is known for the distribution of contacts across age classes [80], our approach still accounts for the major role of children vs. adults in the spread of the disease. Moreover, the validation analysis shows that considering children and adults and the associated mixing and travel behavior is enough to reproduce the spatio-temporal unfolding of the epidemic to a good accuracy. Also, we did not distinguish between symptomatic and asymptomatic infections. Santermans et al. [87] investigated the importance of dealing with symptomatic and asymptomatic infections in an epidemic setting based on differences in mixing patterns between ill and healthy (as a proxy for asymptomatic) individuals. Future research should focus on combining the work of [87] with the study outlined here.

The study is focused on one season only, the 2008/2009 influenza season. Additional seasons may clearly be included in the analysis, however our choice aimed at discounting season-specific effects to avoid uncertainties and discordance found in previous works. Also, we argue that the main effect behind the observed impact is in the interplay between the incidence profile and holidays timing, all other aspects being equal. To fully assess this aspect, we systematically explored earlier and later epidemics than the 2008/2009 season, thus synthetically accounting for other (similar) influenza seasons. We did not consider age-specific susceptibility, as it was largely addressed for example in studies related to the A(H1N1)v2009 pandemic [21]. It would be interesting to explore its effects in future work in addition to social mixing and mobility, thus investigating additional seasonal influenza profiles.

Our experimental scenarios in *travel changes* and *mixing changes* models consider neglecting travel changes and mixing changes in an independent way. The two aspects are expected to be intrinsically dependent, however no study has yet quantified this dependency that could inform a better experimental design. Also, we did not take into account uncertainties associated to the social contact rates estimated from the survey data, as previous work showed their limited impact in fitting serological data [55].

Mobility changes from commuting during regular weekdays to non-regular travel during weekends is obtained from travel statistics. We lack however specific data on travel behavior for adults during school holidays. We therefore assumed that adults would continue commuting during holiday weekdays. While we expect that a fraction of adults would stop commuting at least for few days during breaks as they take time off work, we expect this change in travel fluxes (compensated by additional trips to visit families [45]) to have a negligible effect on the simulated epidemic. More drastic changes on travel, i.e. fully neglecting travel changes as in the *mixing changes* model, indeed did not alter the resulting epidemic.

## 5 Conclusions

With a data-driven spatial metapopulation model calibrated on the 2008/2009 influenza season in Belgium, we showed that regular school closure considerably slows down influenza epidemics and mitigate their impact on the population, because of changes in social mixing that are empirically measured. This may help the management of epidemics and lessen the pressure on the public health infrastructure. The effect is due to both school holidays and weekend closures, the latter periodically dampening transmission. Variations in travel behavior do not lead instead to visible effects. The observed impact strongly depends on the timing of the school closure, and to a lesser extent on its duration. Christmas holiday is the school break generally playing the most important role in mitigating the epidemic course, though variations are observed depending on the influenza season (e.g. early vs. late epidemic). The addition of one week extension after Christmas holiday may represent an additional strategy to further delay the epidemic peak and mitigate its impact.

## Abbreviations

ILI: influenza-like-illness
SEIR: Susceptible-Exposed-Infectious-Recovered
WLS: Weighted Least Square
GP: general practitioner
CI: confidence interval

## Availability of data and material

Commuting data is available upon request from the Directorate General Statistics and Economic Information (DGSEI) [66]. Demographic data is publicly available from Belgian Statistics [1]. Surveillance data is available upon request from the Belgian Scientific Institute of Public Health. Contact data are reported in the paper.

## Author’s contributions

NH, VC and CP conceived and designed the study. KVK analyzed the contact data. GDL developed the model, performed the numerical simulations, and drafted the first version of the manuscript. GDL and PC performed the statistical analysis and prepared the figures. All authors contributed to the interpretation of the results, edited and approved the final manuscript.

## Acknowledgements

We thank Shweta Bansal for useful discussions on the study. We thank the Belgian Scientific Institute of Public Health for making the surveillance data available to us.

